# Lapatinib protects against epileptic seizures via halting glutathione peroxidase 4-dependent ferroptosis

**DOI:** 10.1101/2020.05.28.120733

**Authors:** Jining Jia, Qin Li, Qianyi Sun, Nan Yang, Kangni Chen, Xixi Yin, Honghao Zhou, Xiaoyuan Mao

**Author notes:** Corresponding to: Prof. Xiaoyuan Mao, Department of Clinical Pharmacology, Xiangya Hospital and Institute of Clinical Pharmacology, Central South University, Changsha 410008, China, or.

## Abstract

**Background and Purpose:** Repetitive epileptic seizures trigger massive neuronal death. Therefore, neuroprotection plays a role in preventing neuronal death and inversely suppresses seizure generation. Additionally, some studies have shown ferroptosis, featured by lipid peroxidation (a dominant form of oxidative stress in the brain), is of paramount importance in epileptic seizures. Lapatinib can play a first-line anti-tumor role by targeting oxidative stress and a recent work illustrates the improvement of encephalomyelitis in rodent models after lapatinib treatment. We hypothesize whether lapatinib can protect against ferroptosis in epileptic seizures via regulating lipid peroxidation.

**Experimental Approach:** The epileptic behavior of the mice was recorded after intracranial injection of KA. Western blot and RT-qPCR were used to detect the protein expression of 4-hydroxynonenal (4-HNE) and glutathione peroxidase 4 (GPX4) and the mRNA expression of prostaglandin endoperoxide synthase 2 (PTGS2) in vivo and in vitro. The level of lipid reactive oxygen species (lipid ROS) in cells pretreated with lapatinib was analyzed by flow cytometry.

**Key Results:** Lapatinib remarkably prevented KA-induced epileptic seizures in mice and ferroptosis was involved in the neuroprotection of lapatinib. Compared with the model group, western blot showed that lapatinib significantly upregulated the levels of GPX4. In the ferroptotic cell death model, lapatinib exerted neuroprotection via up-regulating GPX4. Treatment with Ras-selective lethal small molecule 3 (RSL3), a selective GPX4 inhibitor abrogated its anti-ferroptotic potential.

**Conclusions and Implications:** These results illustrated that lapatinib has neuroprotective potential against KA-triggered epileptic seizures via suppressing GPX4-dependent ferroptosis.

**What is already known:**

- Activation of ferroptosis occurs in epileptic seizures.

**What this study adds:**

- Lapatinib protects brain against epileptic seizures via blocking ferroptosis.
- GPX4-dependent ferroptosis is involved in the neuroprotection of lapatinib.

**What is the clinical significance:**

- Inhibition of ferroptosis by lapatinib represents a potential neuroprotective strategy for epileptic patients.

## 1. INTRODUCTION

Epilepsy represents one of the most common neurological disorders and it is characterized by abnormal discharge of highly synchronized neurons in the brain (Feldman, Lapin, Busch, & Bautista, 2018; Loscher, 2011). It is estimated that approximately 65 million people worldwide suffer from epilepsy, with an annual incidence of 80 per 100,000 population (Moshe, Perucca, Ryvlin, & Tomson, 2015). Epileptic patients usually experience social prejudice, misunderstanding and unsuspected stress in life.

Serious and recurrent epileptic seizures undoubtedly trigger neuronal death (Mao, Zhou, & Jin, 2019b). Epilepsy-induced cell death can lead to progressive loss of hippocampal neurons and spatial memory impairment while preventing neuron death can alleviate seizure-induced brain impairment and slow down the progression of epilepsy (Kotloski, Lynch, Lauersdorf, & Sutula, 2002). Besides, due to the limited self-renewal ability of neurons (Sorrells et al., 2018), neuroprotective strategy is beneficial for preventing neuronal death and protection of the structural and functional integrity. Neuroprotection is regarded as a promising therapy for preventing and treating a plethora of neurological disorders (Acharya, Hattiangady, & Shetty, 2008; Neuhaus, Couch, Hadley, & Buchan, 2017). In particular, the antioxidants resveratrol pretreatment can reduce the damage of hippocampal neurons caused by KA, a common chemical reagents for seizure induction (Q. Wang et al., 2004). Another study suggests that brain-derived neurotrophic factor improves hippocampal neuron survival in a model of status epilepticus (Simonato, Tongiorgi, & Kokaia, 2006). Further, early intervention with neural cell transplantation is also helpful for mitigating epileptic seizures and cognitive dysfunction (Acharya et al., 2008). These findings suggest neuroprotection has great benefits for epilepsy. Nowadays, although nearly thirty types of drugs has been developed for curing epilepsy (Y. Wang & Chen, 2019), they mainly alleviate the symptoms of epileptic seizures and control disease progression (Loscher, Klitgaard, Twyman, & Schmidt, 2013), and the ability to protect the structural and functional integrity of neurons is limited. Therefore, it is necessary to seek new neuroprotective drugs for preventing and treating epilepsy.

Lapatinib is a tyrosine kinase inhibitor for treating breast cancer (Konecny et al., 2006). Recently, it is reported to regulate central nervous system (CNS) function. For instance, it was previously reported that experimental autoimmune encephalomyelitis can be effectively improved by lapatinib (Elliot, Christina & William, 2012). Additionally, owing to low molecular weight and lipophilic properties, lapatinib is considered to penetrate the blood-brain barrier in the CNS (Gori et al., 2014), which also supports the notion that it has good therapeutic potential in the CNS. The current study was designed to investigate whether lapatinib protects against epilepsy and the potential molecular mechanism.

Oxidative stress is a critical factor contributing to neuronal death caused by severe and repetitive seizures (Ferriero, 2005). Previous experimental results have also supported the point that epileptic seizures-mediated oxidative stress activates neuronal death (Liang, Ho, & Patel, 2000). In particular, brain is highly vulnerable to oxidative damage due to its consumption of great amounts of oxygen (about 20%) and low antioxidant capacity (Cobley, Fiorello, & Bailey, 2018). Additionally, lipid peroxidation is likely to be a major type of oxidative stress in the brain as there are large amounts of polyunsaturated fatty acids in the neuronal membrane (Cobley et al., 2018). Recently, a non-apoptotic cell death called ferroptosis has been discovered, which is featured by excessive accumulation of iron-dependent lipid peroxidation (Dixon et al., 2012). Ferroptosis has been reported to be associated with a variety of diseases such as Alzheimer’s disease (AD), Parkinson’s disease (PD), cancer and ischemia-reperfusion injury (Guiney, Adlard, Bush, Finkelstein, & Ayton, 2017; Xie et al., 2016). Interestingly, ferroptosis may also be implicated in the pathological process of epilepsy. Emerging evidence shows that ferrostatin-1 (Fer-1), a specific inhibitor of ferroptosis, alleviates cognitive impairment caused by kainic acid (KA)-induced seizures in rats (Ye et al., 2019) and attenuates seizures in pentylenetetrazole and pilocarpine-treated epileptic mice (Mao, Zhou, & Jin, 2019a). Additionally, baicalein can ameliorate posttraumatic epileptic seizure behavior by inhibiting the ferroptosis process (Li et al., 2019). Whereas the former studies have demonstrated that regulation of oxidative stress contributes to the therapeutic effect of lapatinib (Aird et al., 2012; Ma et al., 2017). Therefore, we speculate that lapatinib can eliminate oxidative damage by reducing lipid peroxidation-mediated ferroptosis and eventually exert neuroprotective effects in epilepsy.

## 2 METHODS

### 2.1 Chemicals and reagents

Erastin, lapatinib, Fer-1, liproxstatin-1 (Lip-1) and Ras-selective lethal small molecule 3 (RSL3) were obtained from Selleck Chemicals (Houston, TX, USA). Glutamate (Glu), propidium iodide (PI), Hoechst 33342, KA and deferoxamine (DFO) were obtained from Sigma-Aldrich (St. Louis, MO, USA). Dulbecco’s modified Eagle’s medium (DMEM), Hank’s Balanced Salt Solution (HBSS) and fetal bovine serum (FBS) were obtained from GIBCO (Grand Island, NY, USA). MDA Assay Kit and GSH and GSSG Assay Kit were purchased from Beyotime (Shanghai, China).

### 2.2 Establishment of KA-induced seizure model

Male C57BL/6J mice (6-8 weeks of age, weighing 18-22 g) were obtained from the Animal Unit of Central South University. All C57BL/6J mice were housed in an indoor environment with a 12 h light/12 h dark cycle, 24±2 □ and other specific laboratory conditions, with free access to food and water. Experimental protocols for all animals were approved by the Ethical Committee of Xiangya Hospital Central South University. After anesthetization by intraperitoneal injection of 10% chloral hydrate (0.35 ml/100 g), the mice were fixed on a stereotactic instrument and stereotactically injected with KA (250 ng/µl) into the hippocampus. KA (1 µl) was injected slowly for 5 min and positioned in the hippocampus (anteroposterior [AP] = 2, mediolateral [ML] = 1.8, dorsoventral [DV] = 2.3 mm). After injection, the needle was left in place for additional 10 min to avoid drug reflux. The mice were randomly divided into six experimental groups: 1) sham operation group that received 1 µl PBS injection (5 animals); 2) mice were pretreated p.o. for 21 days on a twice-daily schedule with 100 mg/kg lapatinib alone before PBS administration (5 animals); 3) model group was injected KA (5 animals); 4) and 5) lapatinib groups were received with 50 mg/kg (5 animals) and 100 mg/kg (5 animals) lapatinib for 21 days before KA treatment, respectively; 6) this group was given i.p. for 14 days with ferroptosis inhibitor (3 mg/kg Fer-1) before KA administration.

### 2.3 Behavioral observation

The behavioral changes of epileptic mice were observed successively for 6 h after injection of KA. The symptoms of epileptic seizures were assessed by a scoring system which is defined by Racine (Racine, 1972). The standards of Racine stages were described as follows: stage 0, no convulsions and other responses; stage 1, facial and whisker rhythmic twitching; stage 2, head bobbing and circling; stage 3, myoclonic and spasm in multiple limbs; stage 4, uncontrolled rearing and falling; stage 5, general tonic-clonic seizures with running and jumping; stage 6, death. If the symptoms of the third or higher stage were observed, the animals were considered to have seizures.

### 2.4 Cell culture

Immortalized mouse hippocampal cell line HT22 was cultivated in high-glucose DMEM (C11995500BT, Gibco, USA) containing 10% FBS (10270-106, Gibco, USA), 100 U/ml penicillin and 100 μg/ml streptomycin that maintained in a 5% CO2 incubator at 37 □.

### 2.5 Cell death assay

HT22 cells were cultured in a 24-well plate with a density of 10%. Different concentrations of lapatinib (S1028, Selleck, USA) were pretreated to cells at various time points, and then HT22 cells were exposed to Glu (G8415, Sigma, USA) or erastin (S7242, Selleck, USA). After incubation of Glu or erastin for 8 h, PI and Hoechst 33342 at the concentration of 5 μg/ml were treated for 10 min. Cell death rate was measured by PI (+)/Hoechst (+).

### 2.6 Real-time quantitative PCR

After drug treatment at the indicated time, total RNA from cells and tissues were extracted using TRIzol reagent (Invitrogen, USA) following the manufacturer’s procedures. One microgram of total RNA for each sample was reversely transcribed into cDNA by a commercial kit (RR047A, Takara Bio, Japan) according to the manufacturer’s protocol. Real-time PCR was carried out using SYBR Green PCR Master Mix (RR091A, Takara, Japan) and the quantitative analysis was performed using LightCycler Roche 480 qPCR instrument. The conditions for PCR were as follows: 30□s hot-start at 95□°C followed by 40 cycles of 5□s at 95□°C, 30□s at 55□°C and 30□s at 72□°C; 30□s melting curve at 95 °C. All samples were tested in triplicate, and expression levels were normalized to β-actin gene expression levels. Primer sequences used for real-time quantitative PCR were displayed as follows: prostaglandin endoperoxide synthase 2 (PTGS2) forward: 5’-GGGAGTCTGGAACA TTGTGAA-3’, reverse: 5’-GTGCACATTGTAAGTAGGTGGACT-3’; β-actin forward: 5’-GTGACGTTGACATCCGTAAAGA-3’, reverse: 5’-GCCGGACTCATC GTACTCC-3’.

### 2.7 Immunofluorescence

HT22 cells were cultivated in 35 mm dishes at a density of 15%. After drug treatments, the cells were fixed with 4% paraformaldehyde for 20 min and infiltrated with 0.2% Triton X-100 for 15 min, followed by blocking in donkey serum for 30 min and incubating overnight at 4 □ with primary antibody (GPX4, rabbit, ab125066, 1:200, Abcam) in PBS. On the second day, after three washes with PBS, cells were incubated with donkey anti-rabbit IgG (A21206, 1:100, Thermo Fisher Scientific, USA). Finally, the cell nuclei were stained with DAPI (S2110, Solarbio, China) for 10 min. Fluorescent results were analyzed under a laser-scanning confocal microscope (Nikon, Japan).

### 2.8 Western blot analysis

After drug treatment, HT22 cells or hippocampal tissues were harvested and lysed in RIPA buffer supplemented with protease and phosphatase inhibitor. Cell and tissue lysates were sonicated for 30 sec. The supernatants were retained and subsequently quantified using BCA protein assay kit (P0006, Beyotime Biotechnology, China). In brief, 20 μg proteins for each sample were subjected to sodium dodecyl sulfate–polyacrylamide gel electrophoresis and then electrophoretically transferred to polyvinylidene fluoride membranes. After blocking with TBST containing 5% nonfat milk for 1 h, the membranes were incubated with GPX4 (rabbit, 22 kDa, ab125066, 1:5000, Abcam), 4-hydroxynonenal (4-HNE) (rabbit, 17–76 kDa, ab46545, 1:2500, Abcam), acyl-CoA synthetase long-chain family member 4 (ACSL4) (mouse, 75 kDa, sc-365230, 1:1000, Santa Cruz), solute carrier family 7 member 11 (SLC7A11) (rabbit, 55 kDa, ab175186, 1:5000, Abcam), Anti-5 Lipoxygenase (5-LOX) (rabbit, 78 kDa, ab169755, 1:1000, Abcam) and β-actin (mouse, 43 kDa, A5441, 1:5000, Sigma) overnight at 4 □. The next day, following three washes in TBST, the membranes were incubated with horseradish peroxidase (HRP)-conjugated anti-rabbit IgG (rabbit, A9169, 1:10000, Sigma) or anti-mouse IgG (mouse, A9044, 1:10000,Sigma) at room temperature for 1 h. Immunoreactivity was assessed using the ChemiDoc XRS+ imaging system (Bio-Rad, USA). The protein band intensity was quantified by using Image J software (NIH, USA). The protein expression levels were normalized to β-actin.

### 2.9 Measurement of lipid reactive oxygen species (lipid ROS)

HT22 cells were seeded in 6 well plates at the density of 15% and treated at indicated time points. Cells were washed twice with PBS, trypsinized, and then incubated with 500 μl HBSS (C14175500BT, Gibco, USA) supplemented with 2 μM C11-BODIPY 581/591 (D3861, Thermo Fisher, USA) at 37 □ for 15 min. Then, the cells were washed twice with 500 μl HBSS, centrifuged for 5 min at 3000×*g* and re-suspended in 200 μl PBS. Finally, the fluorescence intensities were detected by flow cytometry and the data were analyzed using FlowJo software.

### 2.10 Measurement of malondialdehyde (MDA) level

The degree of lipid peroxidation can be analyzed by quantification of MDA. MDA content was tested using a specific colorimetric kit (S0131, Beyotime Biotechnology, China) according to the manufacturer’s instructions. Briefly, MDA can react with thiobarbituric acid (TBA) to generate the MDA-TBA adduct which can be measured spectrophotometrically at 532 nm.

### 2.11 Measurement of glutathione (GSH) level

Assessment of GSH level was detected using a GSH and GSSG assay kit (S0053, Beyotime Biotechnology, China) according to the manufacturer’s protocol. The absorbance was assayed at 412 nm with a microplate reader.

### 2.12 Statistical analysis

Results were presented as the mean ± SD. Statistical significance was evaluated by one-way analysis of ANOVA with the Bonferroni test among three groups using GraphPad Prism 5.0 software. Statistical differences with p values less than 0.05 were considered significant.

## 3 RESULTS

### 3.1 Lapatinib protected mice against KA-induced seizures

Initially, we examined whether lapatinib could exert neuroprotective effect and improve seizure behavior in an acute seizure animal model induced by KA. Figure 1a showed the experimental timeline. We established a KA-induced epileptic seizure model by injecting 1 μl 250 ng/μl KA into the hippocampus of mice. The successful establishment of epileptic seizures was assessed by the appearance of obvious symptoms such as rollover and generalized ankylosing seizure. After treatment with lapatinib and KA, seizure behavior was recorded within 6 h (Figure 1b). As shown in Figure 1c-e, pretreatment with lapatinib caused lower seizure score, shorter seizure duration and smaller number of seizures within 90 min in a mouse model of KA-induced epileptic seizures. The above parameters in KA injection group were significantly higher than that in sham operation group (Lu, Zhou, Yang, Deng, & Liu, 2019). Consistently, our preliminary experiments showed similar results (data not shown). Besides, 100 mg/kg lapatinib had no evident toxicity in sham-operated mice (data not shown). And lapatinib had no significant effect on weight gain during the time range of therapy, indicating a good tolerance of lapatinib (Figure 1f). Taken together, these results indicated that lapatinib had a neuroprotective effect in epileptic seizures.

**FIGURE 1.**
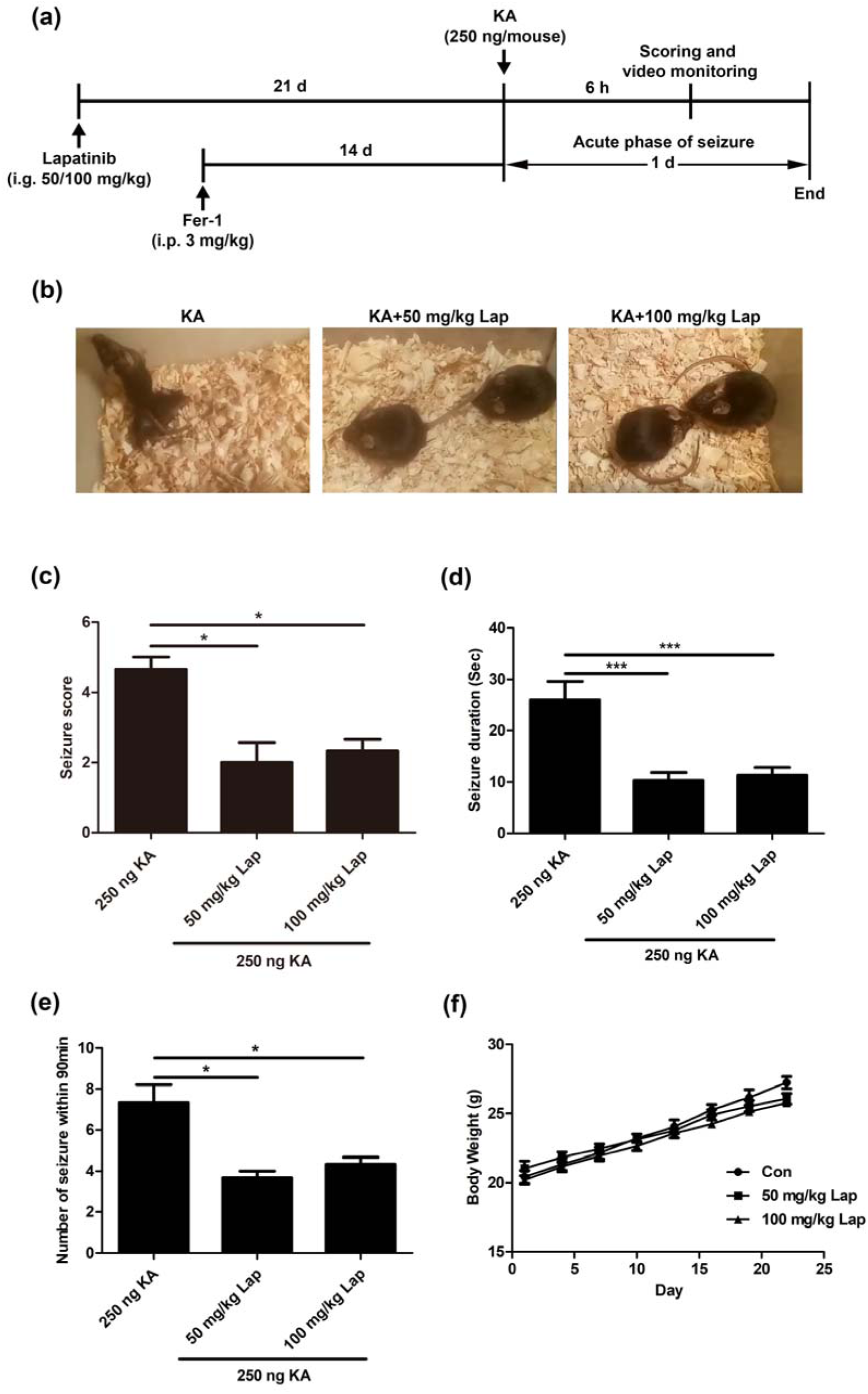
Lapatinib protected mice against KA-induced seizures. (a) Experimental design. (b) Representative images from different groups after pretreatment with lapatinib in KA-induced epileptic seizures. (c-e) Effects of lapatinib on seizure score, number of seizures and average seizure duration. (f) The changes of body weight were recorded during the medication. All results were shown as the mean ± SEM (n = 5), *p < 0.05 and ***p < 0.001.

### 3.2 Lapatinib suppressed ferroptosis in KA-induced epileptic mice

To further investigate the potential molecular mechanism by which lapatinib exerted neuroprotection, the alterations of ferroptosis-related indices including 4-HNE (a common by-product of lipid peroxidation) (Hu et al., 2020) and PTGS2 (a potential molecular marker of ferroptosis) (Jiang et al., 2015) were detected in our present investigation. As shown in Figure 2a,b, KA injection resulted in the upregulation of 4-HNE levels and this phenomenon was reversed after pretreatment with lapatinib and Fer-1 (a selective ferroptosis inhibitor) (Dixon et al., 2012). The mRNA expression of PTGS2 was also dramatically reduced in the seizure model after pretreatment with lapatinib or Fer-1 compared with the KA treatment alone (Figure 2c). Compared with sham operation group, the level of 4-HNE and PTGS2 mRNA were increased in seizure models (Mao et al., 2019a). In preliminary studies, KA-induced mice also exhibited elevations of 4-HNE level and PTGS2 mRNA (data not shown). Collectively, these results indicate that lapatinib’s neuroprotection may be closely related to inhibition of ferroptosis.

**FIGURE 2.**
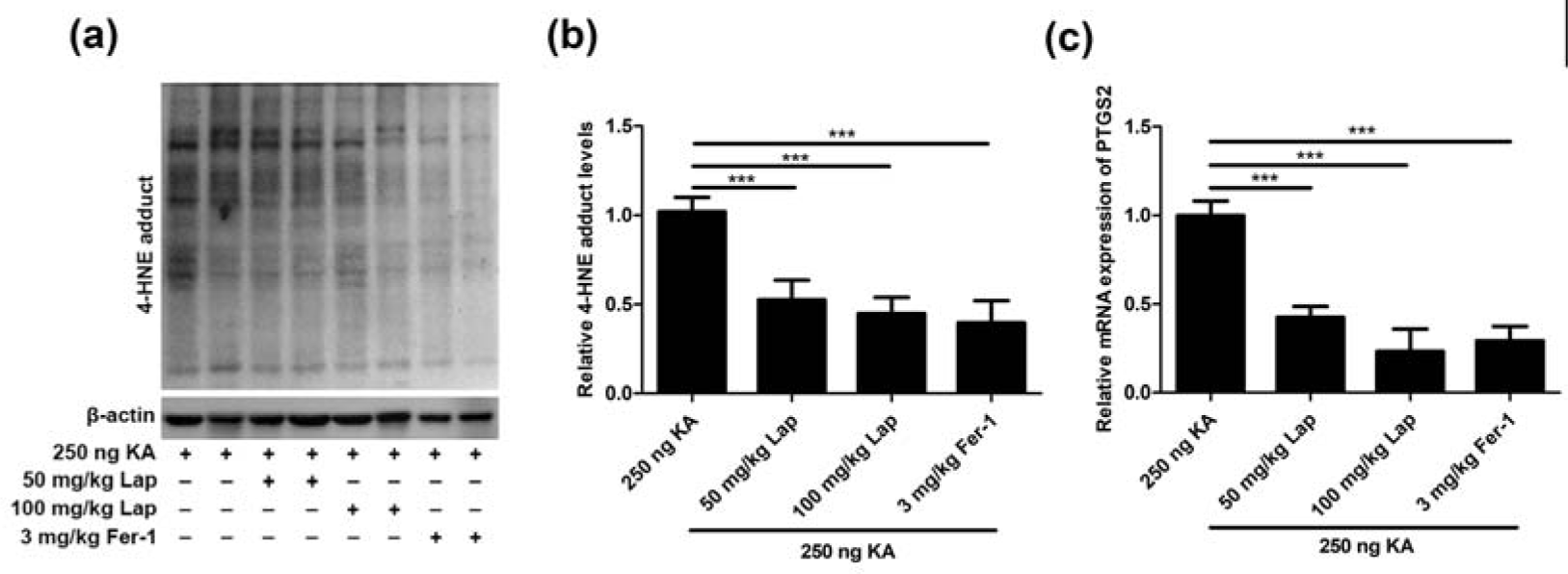
Lapatinib suppressed ferroptosis in KA-induced epileptic mice. (a, b) Western blot showing expression of 4-HNE in hippocampus tissues of KA-induced epileptic mice with different doses of lapatinib and Fer-1 pretreatment. (c) RT-qPCR analysis of PTGS2 mRNA expression pretreated with lapatinib and Fer-1 in epileptic mice of KA injection. All results were shown as the mean ± SEM (n = 5), ***p < 0.001.

### 3.3 Lapatinib exerts neuroprotection against Glu- or erastin-induced cell death in HT22 neurons

To further explore the mechanism on how ferroptosis affects lapatinib’s neuroprotection against KA-induced epileptic seizures, we established an in vitro neuronal ferroptosis model induced by Glu or erastin (a specific ferroptotic inducer). HT22 neuron was currently selected as this cell line was featured by deficient N-methyl-D-aspartate (NMDA) receptor (Jin et al., 2014).

Glu-induced cell damage is an ideal model for studying Glu oxidative toxicity (Jin et al., 2014). Besides, ferroptosis is involved in Glu-induced cell damage as a major type of cell death (Kang, Tiziani, Park, Kaul, & Paternostro, 2014). Simultaneously, we also used the specific ferroptosis model in HT22 cells induced by erastin as a positive control. Next, we detected the effects of lapatinib on ferroptotic cell death in HT22 cells at different doses for different time points. Pretreatment with lapatinib at indicated range of concentrations (1.25-10 μM) significantly diminished HT22 cell death induced by Glu or erastin, and its protective effects reached the peak at 10 μM (Figure 3a,b). Therefore, 10 μM was selected in the following investigations. In order to ascertain the optimal time point, lapatinib at the concentration of 10 μM was treated in the following conditions: pretreatment for 6 h (pre 6 h), pretreatment for 2 h (pre 2 h), co-treatment with Glu or erastin (pre 0 h) and posttreatment for 2 h (post 2 h). It was noteworthy that pretreatment of 10 μM lapatinib for 2 h exerted the most protective effect on Glu- or erastin-induced ferroptosis (Figure 3c,d). And treatment with lapatinib alone had no toxic effect on HT22 neurons (Figure 3e). These findings reveal neuroprotection of lapatinib against Glu-induced neuronal death.

**FIGURE 3.**
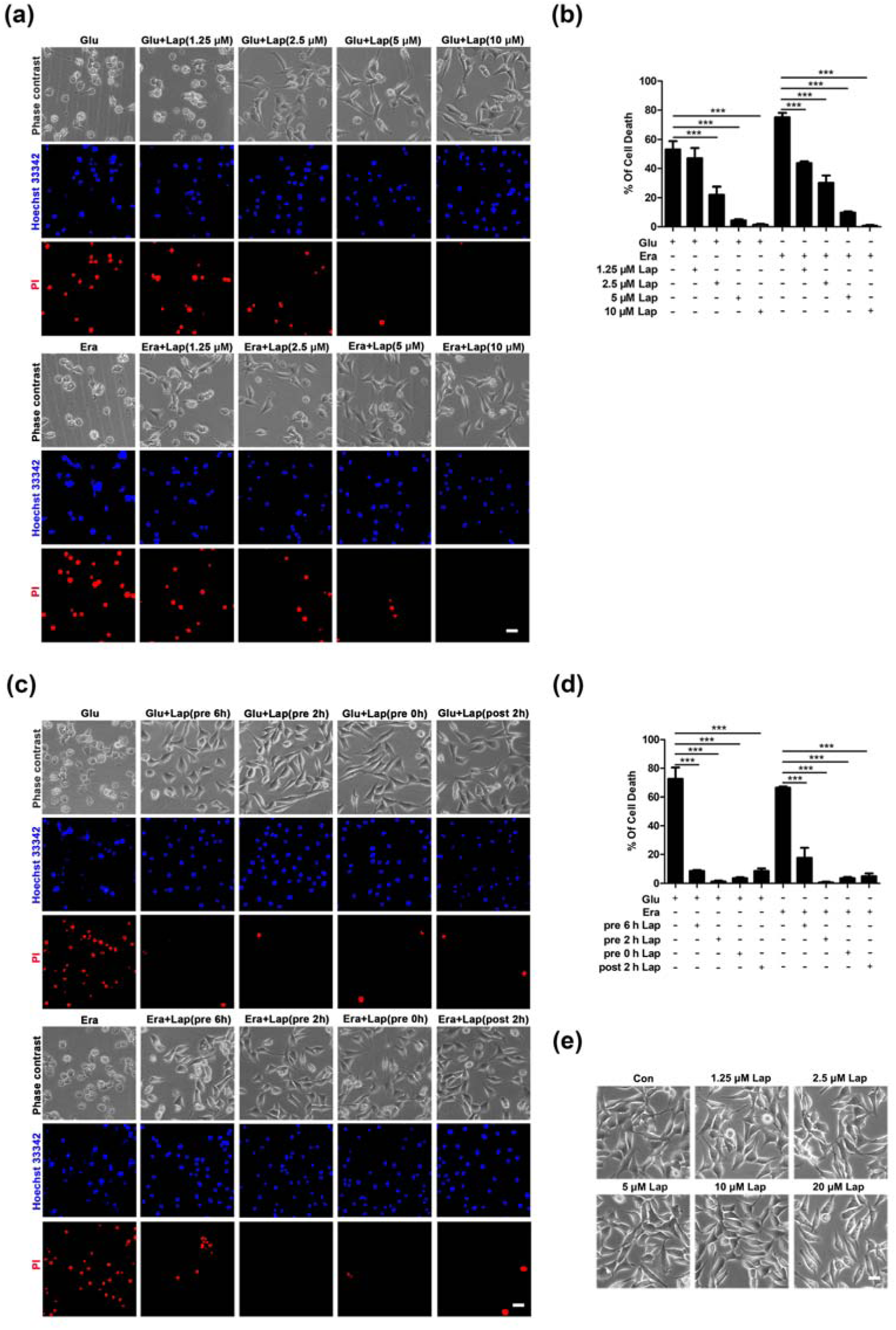
Lapatinib exerts neuroprotection against Glu- or erastin-induced cell death in HT22 neurons. (a) Indicated HT22 cells were treated with Glu (5 mM) or erastin (0.5 μM) for 8 h. Cell morphology was observed by phase-contrast microscopy after pretreatment with lapatinib at different concentrations (1.25, 2.5, 5 and 10 μM). Scale bar: 200 μm. (b) HT22 cells were pre-incubated with lapatinib (1.25, 2.5, 5 and 10 μM) for 2 h, followed with Glu or erastin treatment. Cell death rate was measured by PI (+)/Hoechst (+). (c, d) Lapatinib (10 μM) was put into the neuron cultures for different periods of time (before 6 h, 2 h, simultaneously or post 2 h) of Glu or erastin treatment. The percentage of cell death rate was measured by PI (+)/Hoechst (+). Scale bar: 200 μm. (e) Effects of different concentrations of lapatinib on HT22 cells. Scale bar: 200 μm. All results were shown as the mean ± SEM from three independent experiments, ***p < 0.001.

### 3.4 Lapatinib prevented Glu- or erastin-induced neuronal death possibly by suppressing ferroptosis

In order to further explore whether the neuroprotective effect of lapatinib was correlated with the inhibition of ferroptosis, lipid peroxidation (the feature of ferroptosis) was detected in our present work. As shown in Figure 4a,b, an evident decrease of lipid ROS accumulation was observed in Glu- or erastin-induced ferroptosis after lapatinib pretreatment. In ferroptotic cell death model in HT22, lipid ROS content was increased significantly compared with the control group (Jelinek et al., 2018). We detected overwhelming lipid ROS after Glu and erastin exposure in HT22 cells (data not shown). And lapatinib also remarkably suppressed other ferroptotic indices, including 4-HNE levels (Figure 4c), MDA (a product of lipid metabolism) (Figure 4d), and PTGS2 mRNA (Figure 4e). Additionally, the protective effect of lapatinib combined with ferroptosis inhibitor (Fer-1, Lip-1 and DFO) on Glu- or erastin-induced oxidative toxicity was not significantly different from that of lapatinib alone (Figure 4f,g). These experimental data suggest that lapatinib exerts neuroprotective effects possibly by targeting ferroptosis.

**FIGURE 4.**
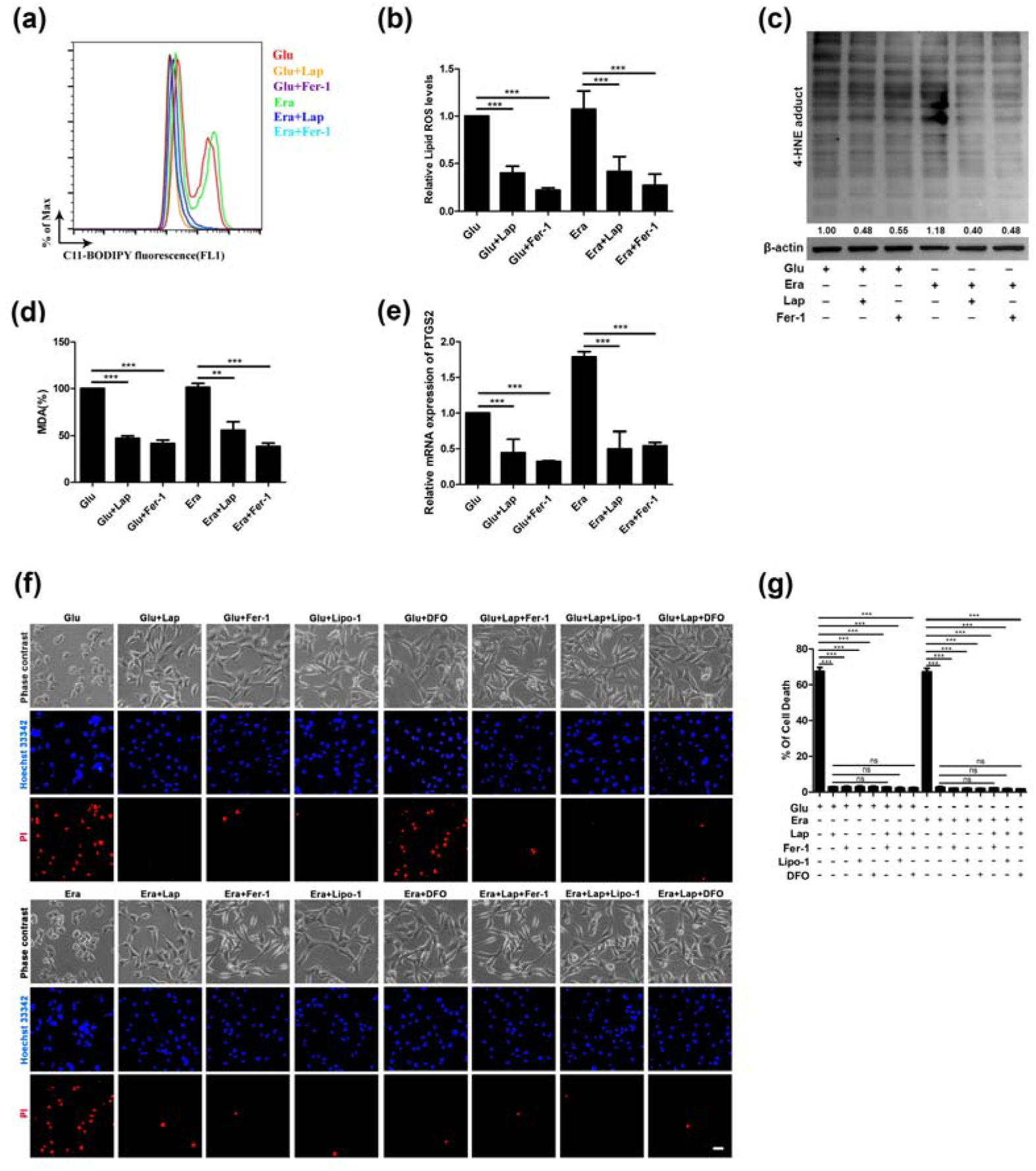
Lapatinib prevented Glu- or erastin-induced neuronal death possibly by suppressing ferroptosis. (a-d) Detection of lipid ROS, 4-HNE and MDA content in the Glu- or erastin-induced HT22 cell injury model following lapatinib (10 μM) and Fer-1 (12.5 μM) pretreatment for 2 h. (e) RT-qPCR analysis of PTGS2 mRNA expression pretreated with or without lapatinib (10 μM) and Fer-1 (12.5 μM) in HT22 cells induced by Glu or erastin. (f, g) To observe and compare the effect between lapatinib and ferroptosis inhibitors co-intervention group and lapatinib intervention alone group on the Glu- or erastin-induced cell injury model. Scale bar: 200 μm. All results were shown as the mean ± SEM from three independent experiments, ns means the difference is not statistically significant, **p < 0.01, and ***p < 0.001.

### 3.5 Lapatinib inhibited neuronal ferroptosis by upregulating GPX4

Next, we explored the molecular mechanism of lapatinib’s neuroprotection against neuronal ferroptosis. We examined diverse factors including GPX4 (Ingold et al., 2018), SLC7A11 (Sehm et al., 2016), 5-LOX (Sun, Zhou, & Mao, 2019) and ACSL4 (Doll et al., 2017) (Figure 5a) that were associated with lipid peroxidation.

**FIGURE 5.**
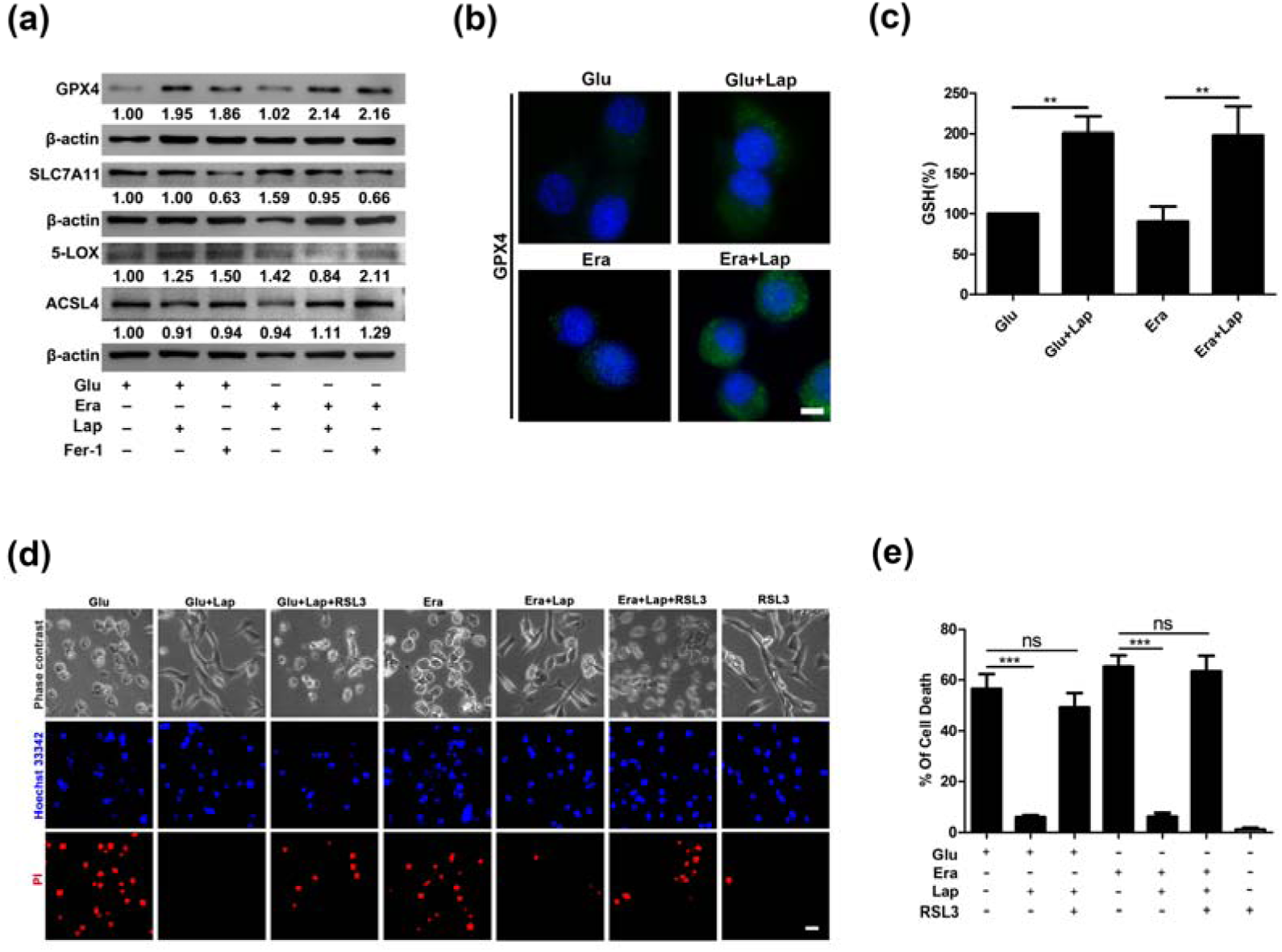
Lapatinib inhibited neuronal ferroptosis by upregulating GPX4. (a) Western blot analysis of ferroptosis-related proteins including GPX4, SLC7A11, 5-LOX and ACSL4 pretreated with lapatinib (10 μM) and Fer-1 (12.5 μM) for 2 h in Glu- or erastin-induced HT22 cell injury model. (b) Immunofluorescence analysis of GPX4 expression after pretreatment for 2 h with lapatinib (10 μM), followed by exposure to Glu or erastin for another 8 h in HT22 cells. Scale bar: 50μm. (c) The level of GSH was assessed following lapatinib (10 μM) pretreatment for 2 h. (d, e) To observe the effect between lapatinib (10 μM) and RSL3 (1 nM) co-intervention group and lapatinib intervention alone group on the Glu- or erastin-induced cell injury model. Scale bar: 200 μm. All results were shown as the mean ± SEM from three independent experiments, ns means the difference is not statistically significant, **p < 0.01, and ***p < 0.001.

We found the most evident upregulation of GPX4 in the Glu- or erastin-induced HT22 cell injury model after pretreatment with lapatinib. Meanwhile, we also detected the expression level of GPX4 in the Glu- or erastin-induced cell damage model by immunofluorescence. Compared with the model group, the immunofluorescence signal of GPX4 was significantly upregulated in the lapatinib pretreatment group (Figure 5b). Additionally, GPX4 transforms toxic lipid peroxide into nontoxic lipid alcohols by using GSH as an auxiliary factor in lipid peroxidation-dependent ferroptosis (Imai, Matsuoka, Kumagai, Sakamoto, & Koumura, 2017). As shown in Figure 5c, in HT22 cells following Glu or erastin treatment, lapatinib increased intracellular GSH levels, which inversely correlate with lipid peroxidation (Ayala, Munoz, & Arguelles, 2014). GSH is the substrate for GPX4 to exert an antioxidant effect. Consistent with previous reports, cellular GSH levels were significantly decreased in HT22 cells after exposure to Glu or erastin (Liu et al., 2015). Given that consumption of GPX4 in vitro can be reversed by lapatinib, we subsequently explored whether treatment with RSL3, a specific GPX4 inhibitor could reverse the protective effect of lapatinib. It was obvious that intervention of RSL3 abrogated the improvement of lapatinib on neuronal survival (Figure 5d,e). Overall, these data strongly document that lapatinib exerts an inhibitory role on neuronal ferroptosis by upregulating the expression of GPX4.

### 3.6 GPX4 was involved in the inhibition of ferroptosis of lapatinib in KA-epileptic seizures

Subsequently, we validated the above in vitro results in a mouse model of KA-induced epileptic seizures. Consistent with the experimental results in vitro, the expression of GPX4 level in the hippocampus was significantly increased after pretreatment with lapatinib compared to the KA-induced model group (Figure 6a,b). Recent studies have found that decreased expression of GPX4 was detected in the mouse models with epilepsy compared with sham operation group (Li et al., 2019; Mao et al., 2019a). Focus on previously detected molecular targets, quantification of the western blot reflected that lapatinib pretreatment slightly upregulated the level of SLC7A11 (Figure 6a,c), but there were no significant changes in the levels of 5-LOX and ACSL4 (Figure 6a,d,e). The above results further established the role of GPX4. Generally, these results demonstrate that lapatinib protects mice against ferroptosis in KA-triggered epileptic seizures via halting GPX4-dependent ferroptosis.

**FIGURE 6.**
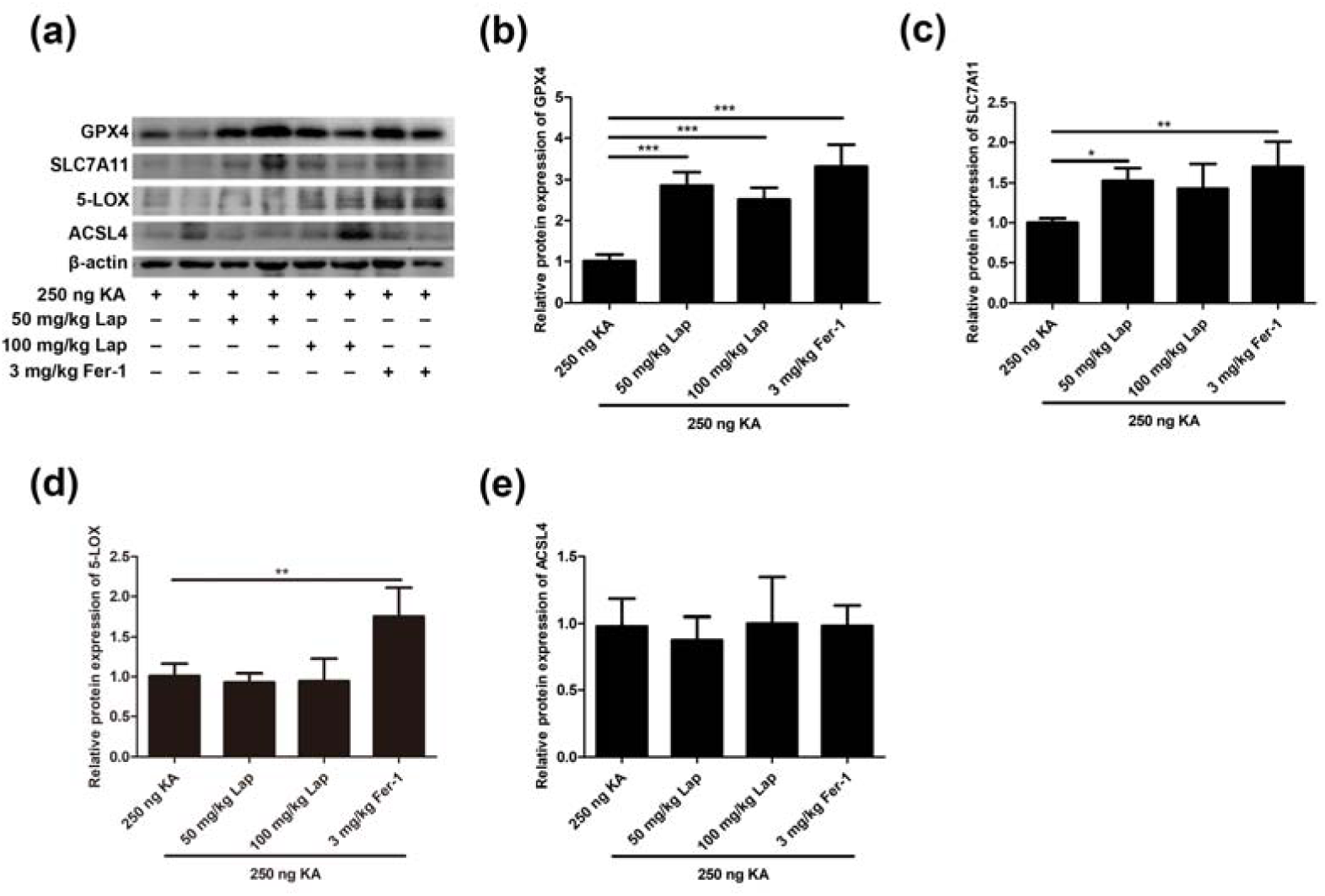
GPX4 was involved in the inhibition of ferroptosis of lapatinib in KA-epileptic seizures. (a) Western blot showing expression of GPX4, SLC7A11, 5-LOX and ACSL4 in hippocampus tissues of KA-induced epileptic mice with different doses of lapatinib and Fer-1 pretreatment. (b-e) Quantitative analysis of GPX4, SLC7A11, 5-LOX and ACSL4. All results were shown as the mean ± SEM (n = 5), *p < 0.05, **p < 0.01 and ***p < 0.001.

## 4 DISCUSSION

The major finding of our current work was that lapatinib exerted neuroprotection against epileptic seizures. And we further found that the neuroprotective potential of lapatinib was at least in part related to the inhibition of neuronal ferroptosis dependent of GPX4.

Recently, drug repurposing (also known as drug repositioning) is regarded as a new strategy of drug development and has been increasingly emphasized (Oprea & Mestres, 2012). Drug repurposing is used to determine the new uses of approved or researched drugs. In order to avoid the long preparation period of new drugs, antiepileptic drugs can be determined by drug repositioning (Brueggeman et al., 2019). Compared with the development of new drugs, the main advantage of the proposed method is that the pharmacokinetic, pharmacodynamic and toxicological properties have been defined by clinical research (Nosengo, 2016). Here, we focus on the tyrosine kinase inhibitors, whose therapeutic characteristics are to achieve the anti-cancer effect through high specificity of biological targeting. A growing number of researches have demonstrated that these tyrosine kinase inhibitors not only are beneficial for antagonizing cancer but also played a critical role in the non-neoplastic field. For example, asthmatic responses and autoimmune arthritis could be effectively cured by imatinib (Berlin & Lukacs, 2005; Paniagua et al., 2006). Focus on therapeutic strategies for neurological diseases, sunitinib, a traditional anti-cancer drug, provided an attractive therapy for protecting HIV neurotoxicity (Wrasidlo et al., 2014). Several studies reported that imatinib also ameliorated AD by inhibition amyloid-β formation (He et al., 2010). Remarkably, lapatinib had therapeutic effect on experimental autoimmune encephalomyelitis (Elliot, Christina & William, 2012). These findings supported the application of lapatinib in the CNS.

Our results showed that lapatinib’s neuroprotection was achieved by suppressing ferroptosis in neurons. Ferroptosis, a newly discovered form of regulated necrosis, is characterized by iron accumulation in cellular and excessive occurrence of lipid peroxidation (Dixon et al., 2012). Numerous reports have established that ferroptosis was highly correlated with multiple physiological and pathological processes, such as cancer, neurotoxicity, acute kidney failure, liver injury and so on (Xie et al., 2016). Here, we focus on ferroptosis in epileptic seizures to expand and discuss. For example, it was found that ferroptosis was closely associated with cognitive impairment and ferroptosis inhibitor Lip-1 could significantly improve behavioral seizures in the iron chloride-induced mouse posttraumatic epilepsy model (Li et al., 2019). Similarly, in the KA- or pilocarpine-induced seizures model, treatment with Fer-1 could remarkably counteract epileptic seizures (Mao et al., 2019a; Ye et al., 2019). Previous investigations supported that lapatinib exerted anti-tumor effect via induction of ferroptosis (Ma et al., 2017; Ma, Henson, Chen, & Gibson, 2016). In our current study, we uncovered that lapatinib protected against epileptic seizures via suppressing ferroptosis in murine models by KA injection. Our researches further proved that GPX4, a vital antioxidant enzyme (Fricker, Tolkovsky, Borutaite, Coleman, & Brown, 2018), was involved in the neuroprotective effects of lapatinib. GPX4 plays a decisive role in inhibiting the accumulation of lipid ROS and scavenges lipid peroxidation under oxidative stress. In contrast, a specific inhibitor of GPX4 can initiate ferroptosis (W. S. Yang et al., 2014). In our previous work, we found that RSL3 at a concentration of 12.5 nM for 8 h exerted potent damage on HT22 cells. However, we chose the sublethal dose of RSL3 (1 nM) to act on HT22 cells. It seems more convincing to reverse the protective effect of lapatinib on the cytotoxic model caused by Glu or erastin, while RSL3 at the concentration of 1 nM does not cause cell death. GPX4 is a major regulator of ferroptosis. And lots of researches have shown that GPX4 is related to the development process of human diseases (Friedmann Angeli et al., 2014; Matsushita et al., 2015). Mice lacking GPX4 in the brain had no resistance to immune dysfunction and tissue impairment. In fact, motor neurons were highly sensitive to GPX4 depletion and accompanied by regulatory cell death with features of ferroptosis (Chen, Hambright, Na, & Ran, 2015). Further studies showed that induction of GPX4 knockout in adult mice led to rapid death with the loss of hippocampal neurons (Yoo et al., 2012). Given the indispensability of GPX4 for neuronal function, selective deletion of GPX4 in the brain contributed to cognitive impairment and neurodegeneration (Hambright, Fonseca, Chen, Na, & Ran, 2017). Besides, the survival of interneurons exclusively depended on selenocysteine-containing GPX4, thereby preventing fatal epileptic seizures (Ingold et al., 2018).

Collectively, our present work showed that lapatinib, a well-known anti-tumor drug, had protective effects against epileptic seizures. And GPX4-mediated ferroptosis was involved in the neuroprotective effect of lapatinib in epileptic seizures (Figure 7). As we all know, seizures have long been considered as a complication of AD, ischemic stroke and other neurological diseases (Camilo & Goldstein, 2004; Larner, 2010). In addition, PD, schizophrenia, and cerebral infarction appear to be primarily associated with cognitive dysfunction (Bennett, Schneider, Bienias, Evans, & Wilson, 2005; Green, 2006; Y. Yang, Tang, & Guo, 2016). So, whether lapatinib can have beneficial effects on other brain diseases except epilepsy remains to be further explored.

**FIGURE 7.**
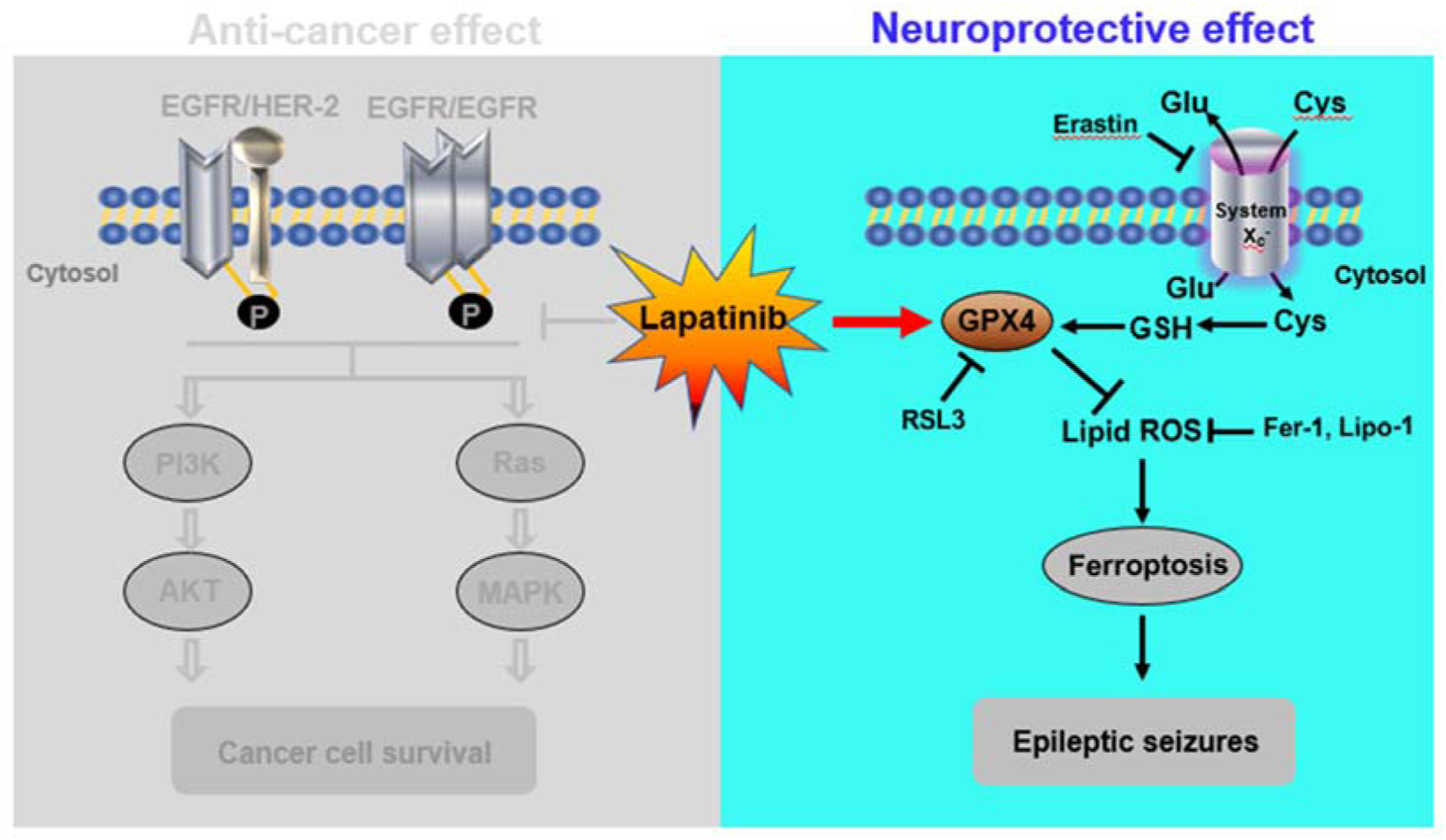
Schematic diagram depicting a novel role of suppressing neuronal ferroptosis for lapatinib in epileptic seizures. The graph not only depicted the traditional anti-cancer effect of lapatinib, but also showed the neuroprotection of lapatinib in KA-induced epileptic seizures in mice. Lapatinib protected against the KA-induced epileptic seizures by inhibiting GPX4-dependent neuronal ferroptosis.

## Abbreviations

KA: kainic acid;
Glu: glutamate;
lipid ROS: lipid reactive oxygen species;
MDA: malonaldehyde;
4-HNE: 4-hydroxynonenal;
GPX4: glutathione peroxidase 4;
CNS: central nervous system;
Fer-1: ferrostatin-1;
Lip-1: liproxstatin-1;
DFO: deferoxamine;
DMEM: Dulbecco’s modified Eagle’s medium;
HBSS: Hank’s Balanced Salt Solution;
FBS: fetal bovine serum;
PI: propidium iodide;
ACSL4: acyl-CoA synthetase long-chain family member 4;
SLC7A11: solute carrier family 7 member 11;
5-LOX: Anti-5 Lipoxygenase;
PTGS2: prostaglandin endoperoxide synthase 2;
GSH: glutathione;
AD: Alzheimer’s disease;
PD: Parkinson’s disease

## AUTHOR CONTRIBUTIONS

X.M. and J.J. conceived and designed the study. J.J. performed the experiments. Q.L., Q.S., X.Y., K.C. and N.Y. provided experimental support. J.J. wrote the paper. X.M. and H.Z. revised the manuscript.

## ACKNOWLEDGEMENTS

This work was financially supported by the National Natural Science Foundation of China (No. 81974502 and 81491293), Natural Science Foundation of Hunan Province (No. 2017JJ3479) and the Fundamental Research Funds for the Central Universities of Central South University (2019zzts788).

